## Supplementary Information for "Thermal phenotypic plasticity of pre- and post-copulatory male harm buffers sexual conflict in wild *Drosophila melanogaster*"

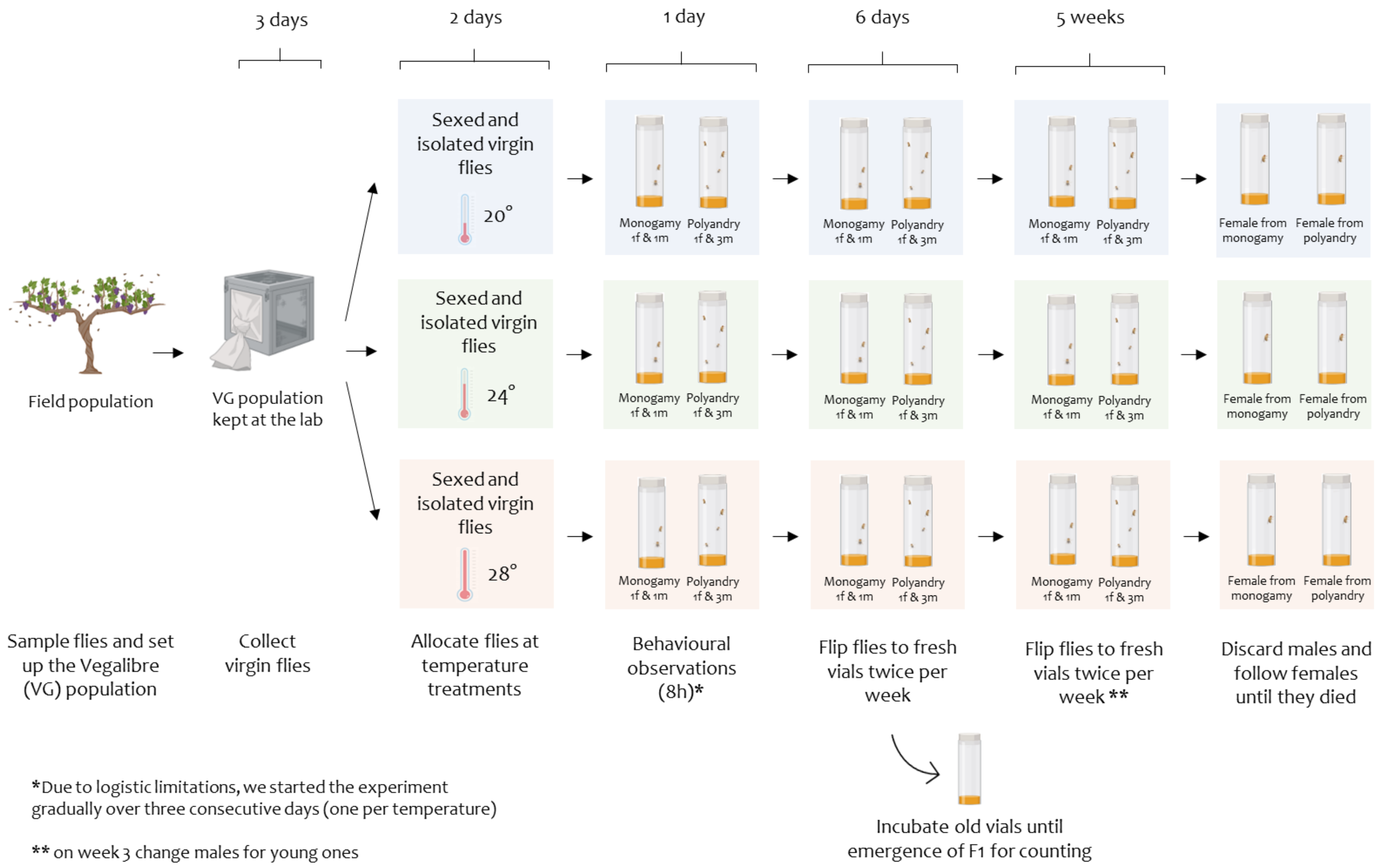


**Figure 1 – figure supplement 1.** Fitness and behavioural assay design (Experiment 1). Referred in main text as S1.1.

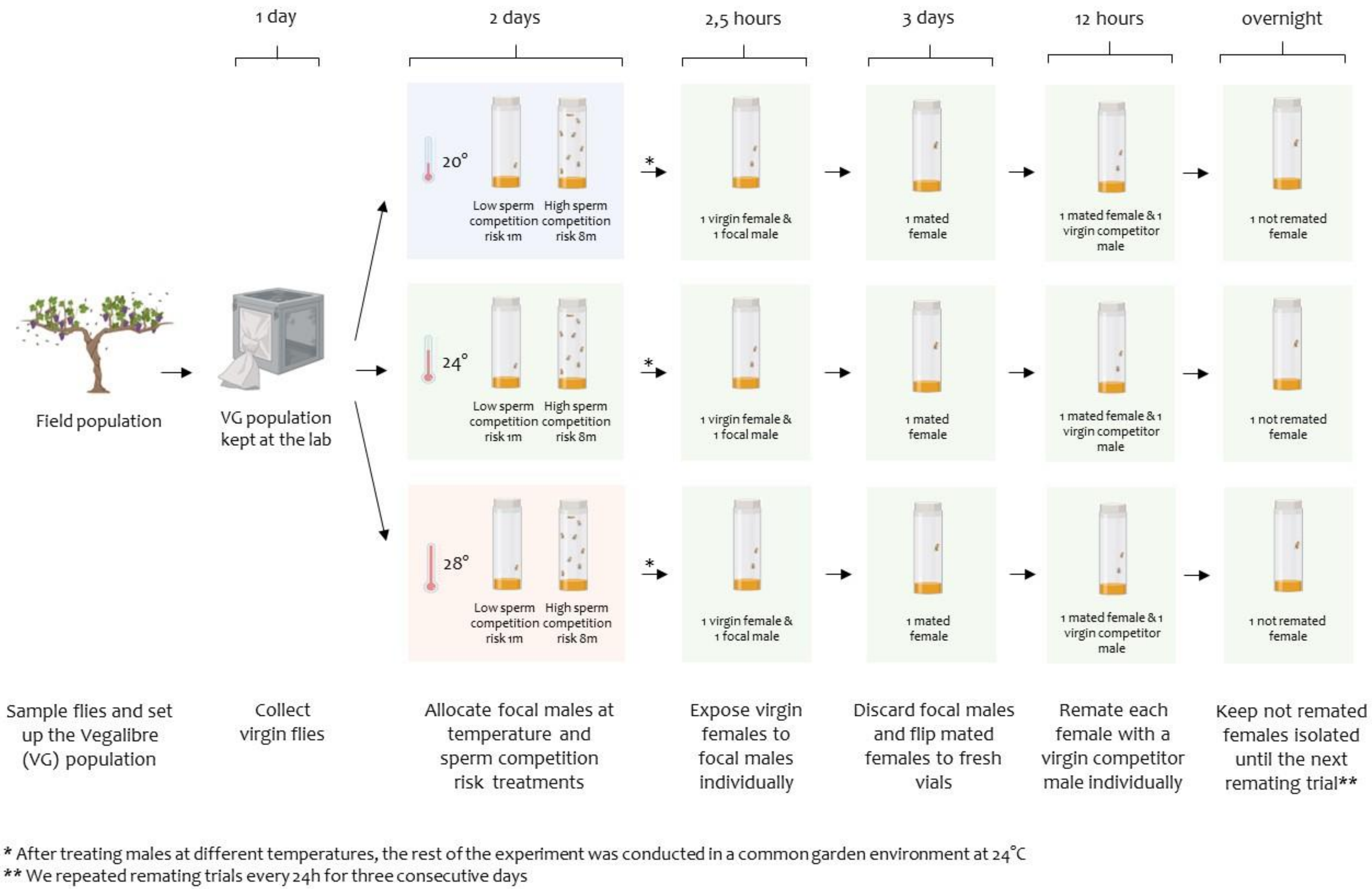


**Figure 1 – figure supplement 2.** Receptivity assay design (Short treatment duration – 48 hours-, experiment 2). Referred in main text as S1.2.

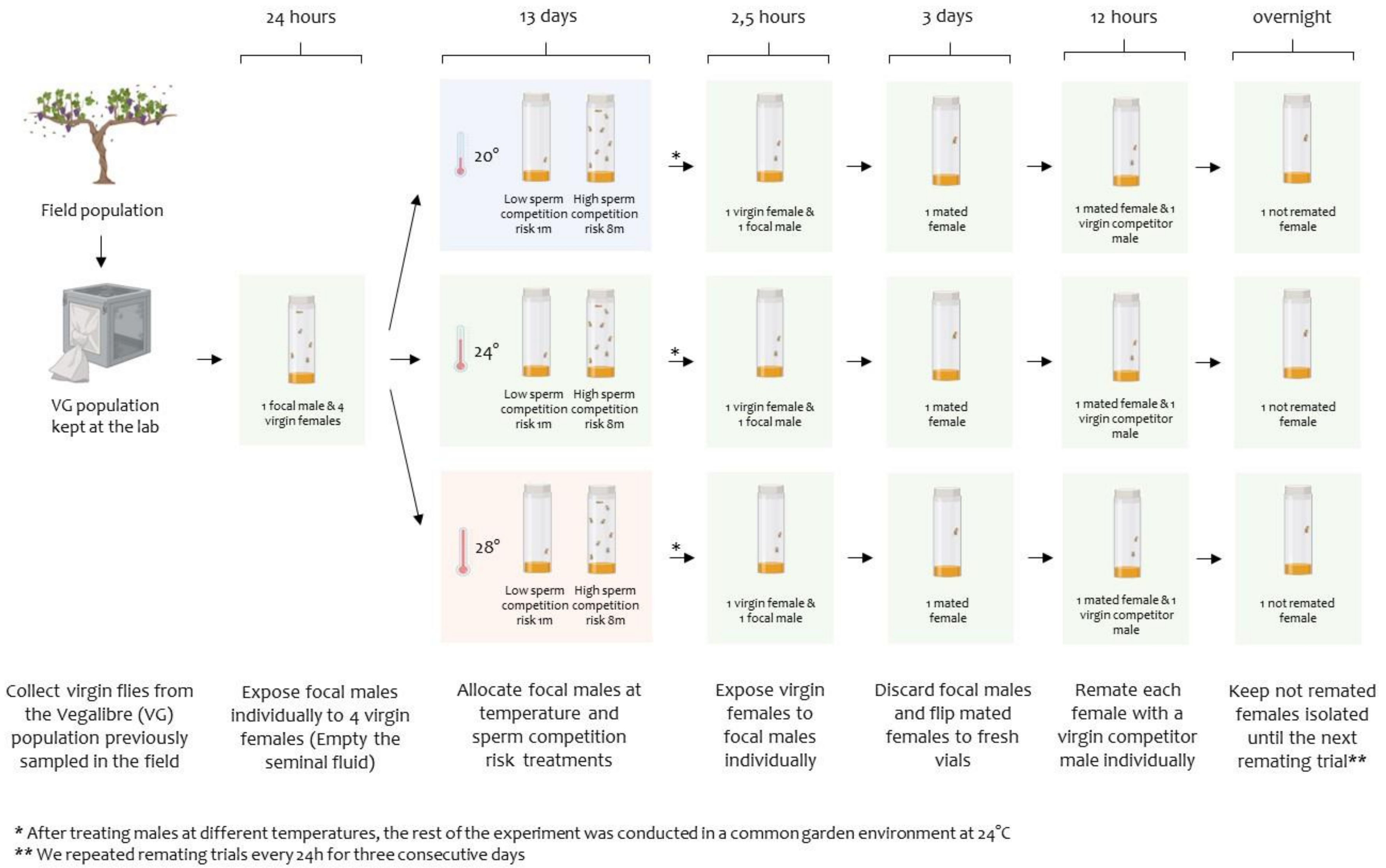


**Figure 1 - figure supplement 3.** Receptivity assay design (Long treatment duration – 13 days-, experiment 3). Referred in main text as S1.3.

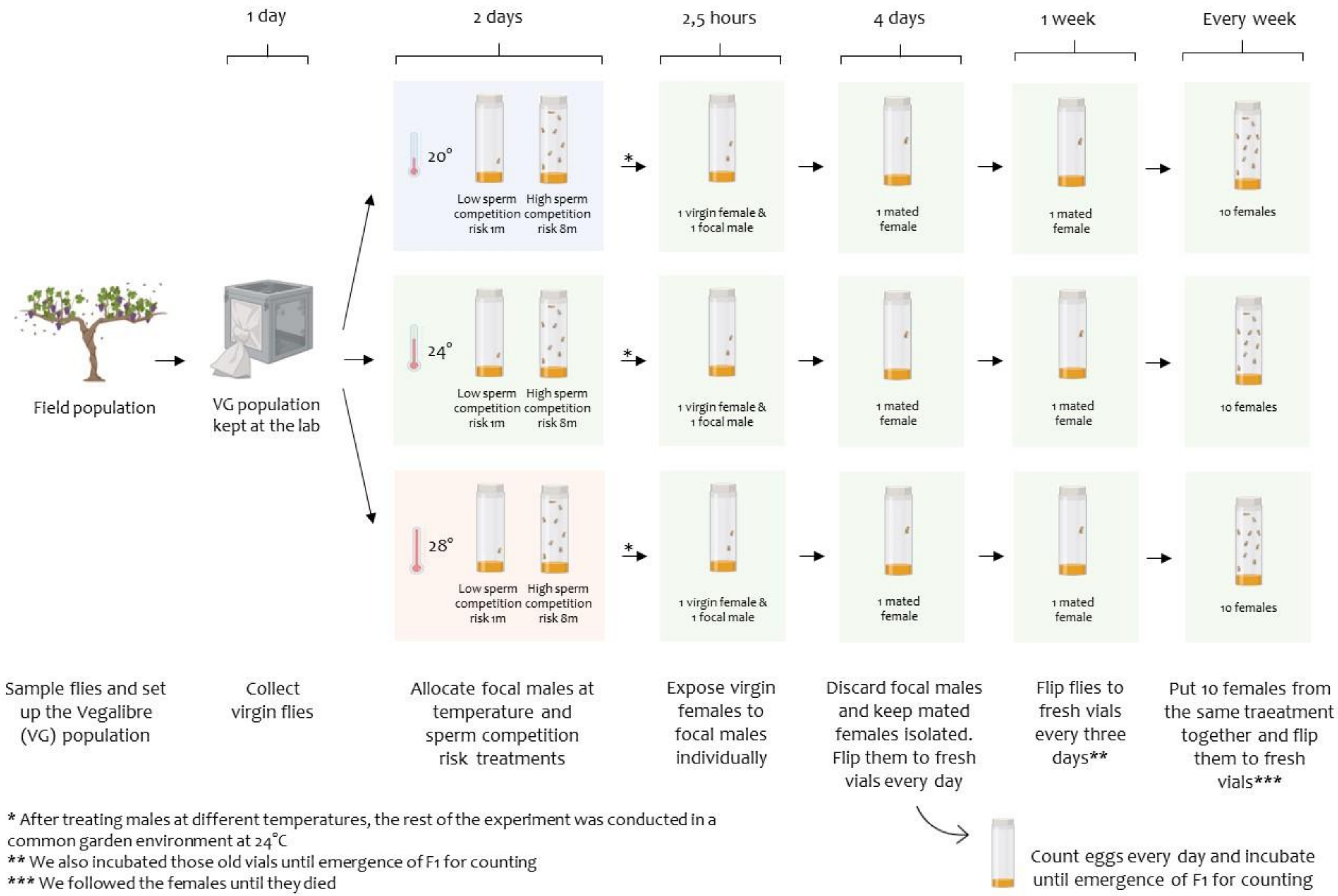


**Figure 1 – figure supplement 4.** Fecundity and survival assay design (Short treatment duration – 48 hours-, experiment 4). Referred in main text as S1.4.

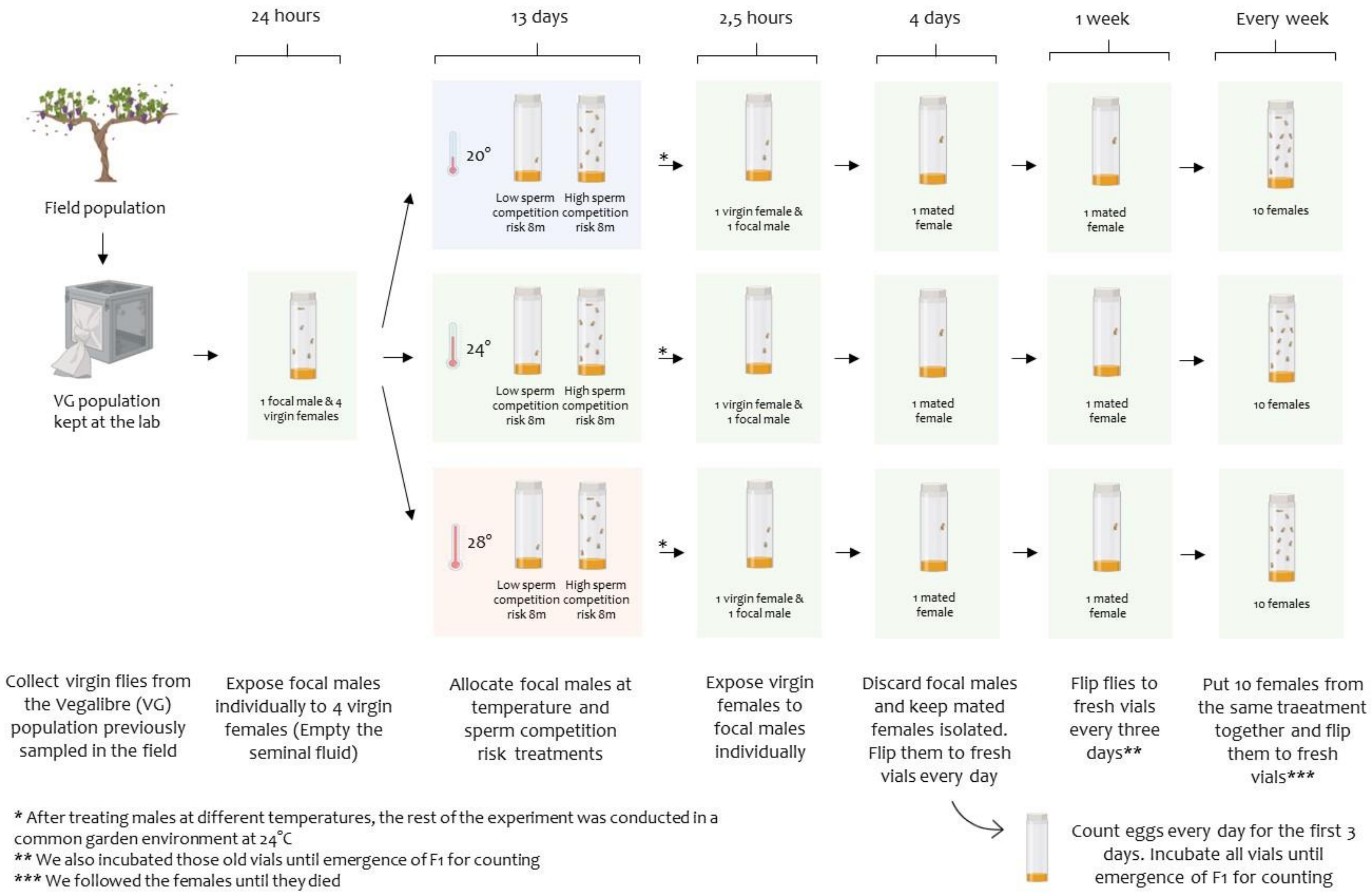


**Figure 1 – figure supplement 5.** Fecundity and survival assay design (Long treatment duration – 13 days-, experiment 5) . Referred in main text as S1.5.

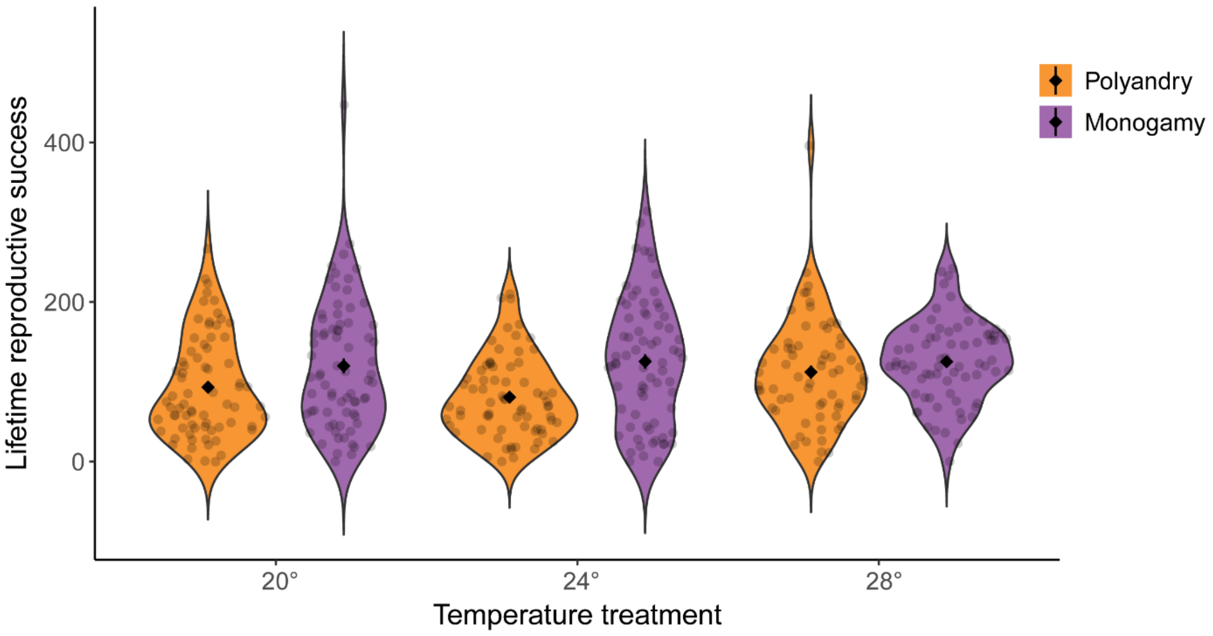


**Figure 2 - figure supplement 1.** Violin plot for female reproductive success across temperature and mating system treatments. Referred in main text as S2.1.

a)

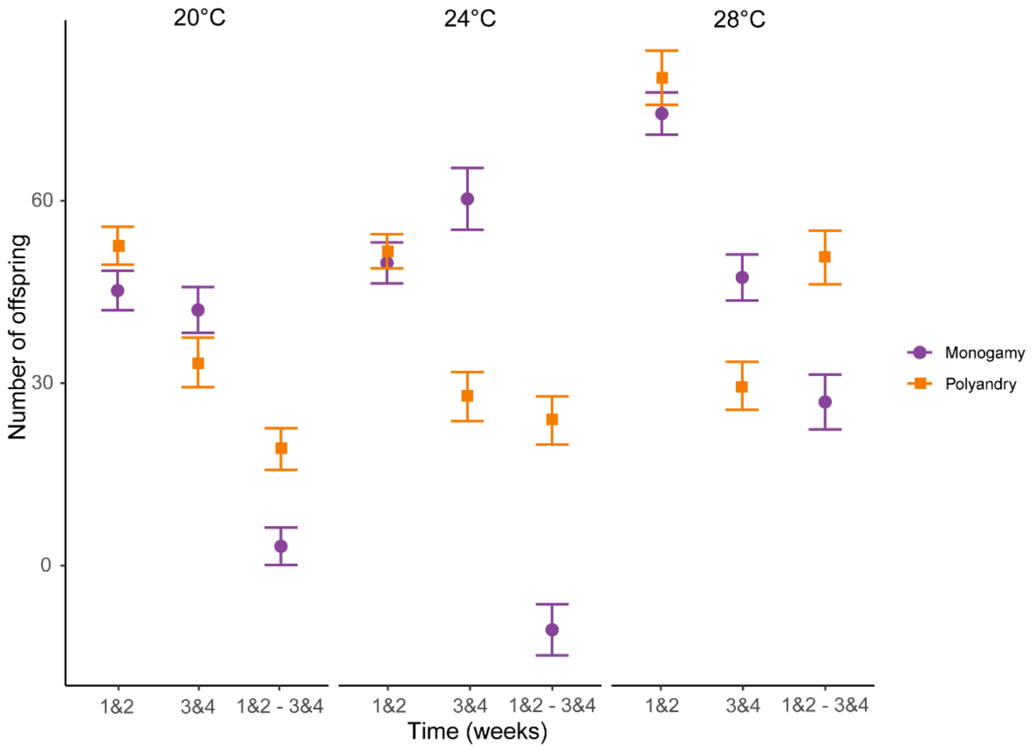


b)

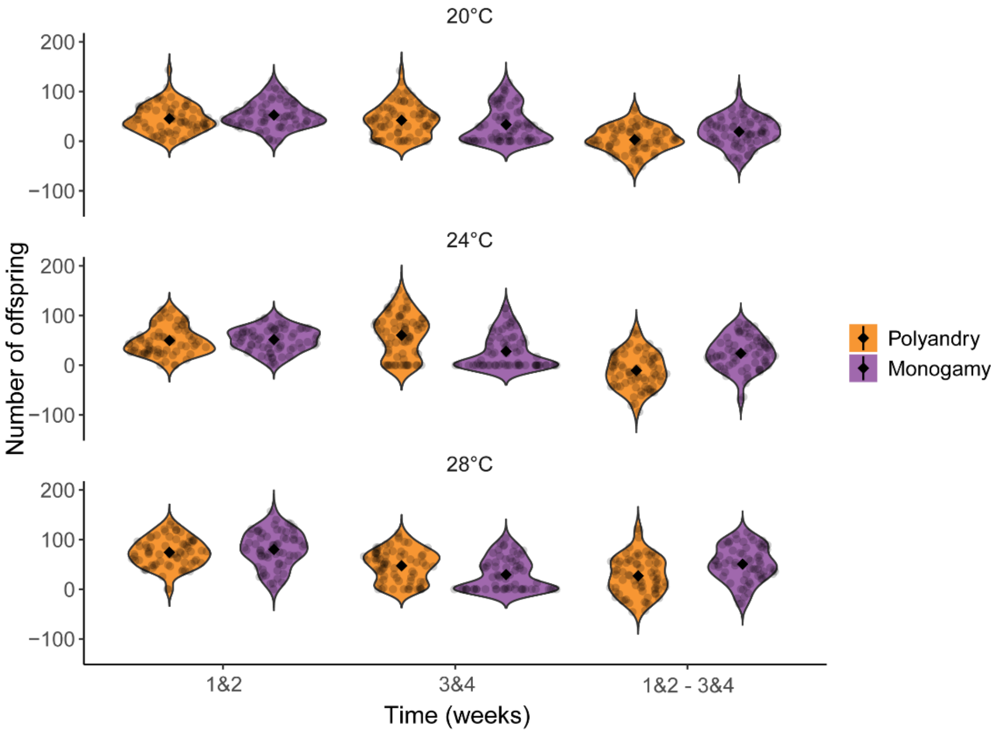


**Figure 2 – figure supplement 2.** Early reproductive rate (number of offspring produced during the first two weeks of age), late reproductive rate (number of offspring produced during the second two weeks of age) and reproductive ageing (number of offspring produced over weeks 1-2 vs. 3-4) plots. a) Mean ± SEM number of offspring in monogamy vs. polyandry mating system treatments, across the different temperature treatments. b) violin plots. Referred in main text as S2.2.

a)

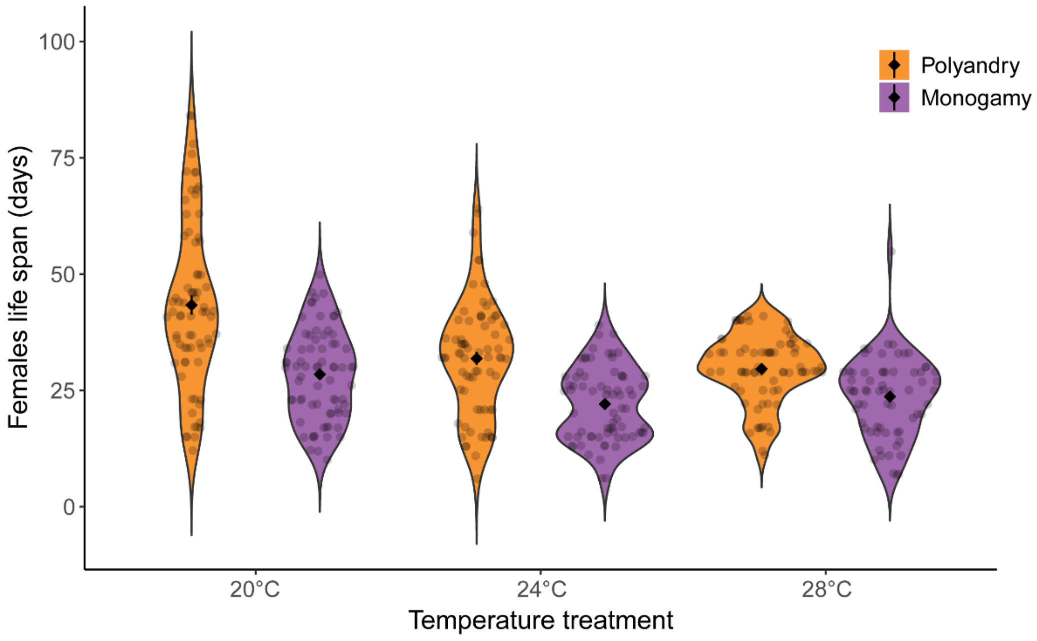


b)

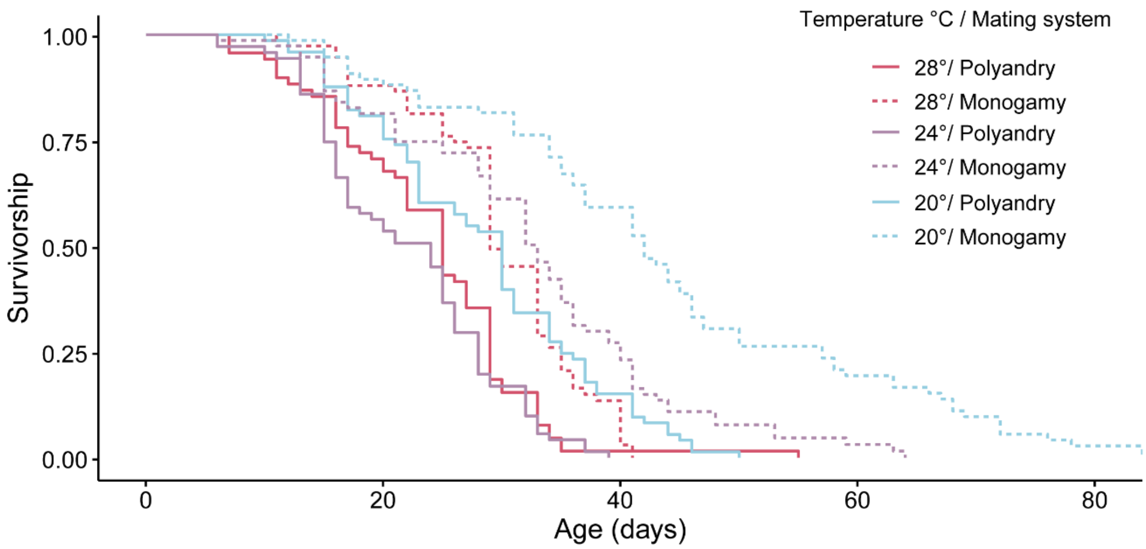


**Figure 4 - figure supplement 1.** a) Violin plot of male harm effect on female lifespan across temperature and mating system treatments. b) Survival plot from the Cox proportional hazard model of male harm effect on female lifespan across temperature and mating system treatments. Referred in main text as S4.1.

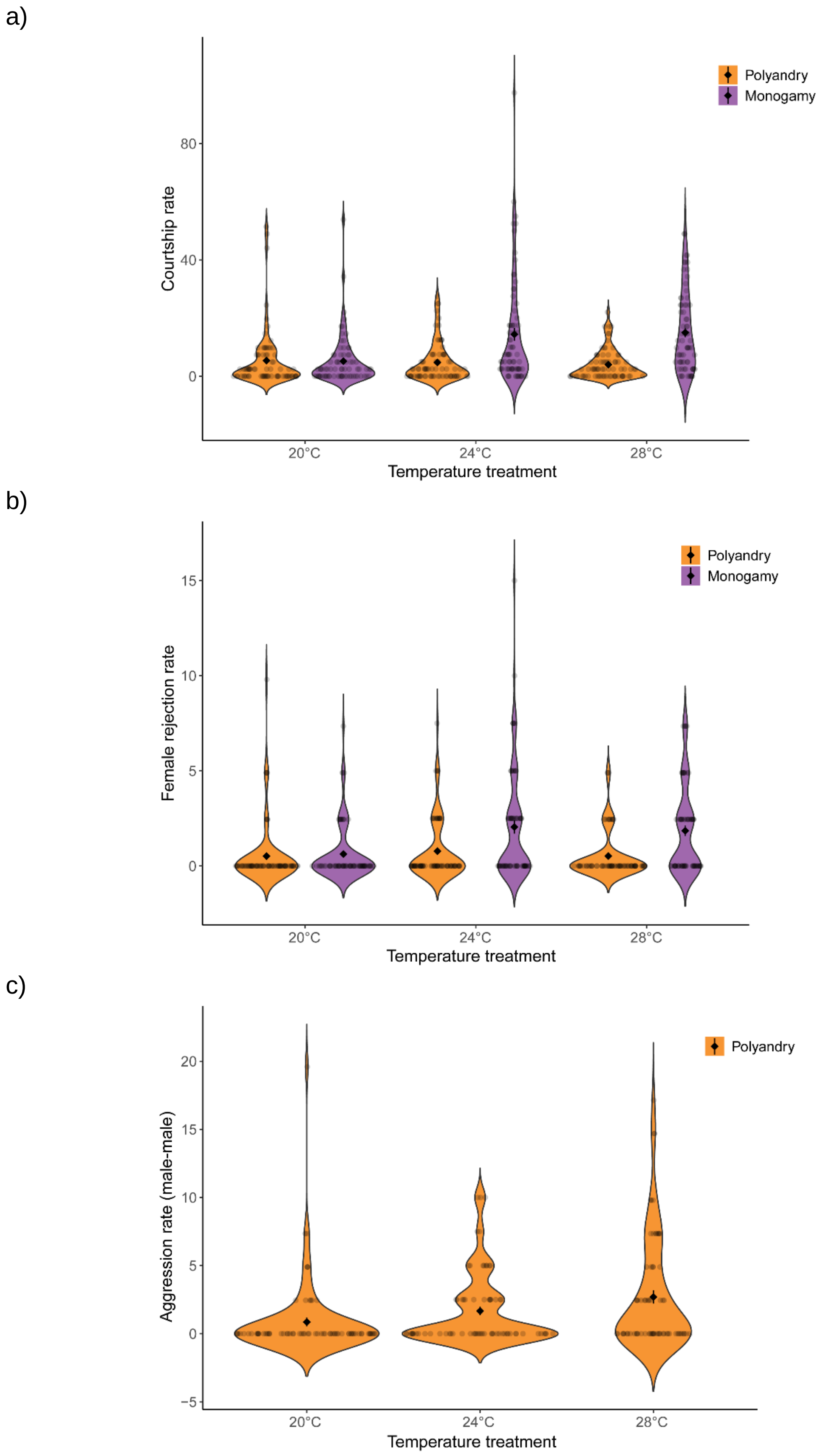


**Figure 5 – figure supplement 1.** Violin plot for male harm effect on a) courtship rate b) rejection rate across temperature and mating system treatments. c) Violin plot for polyandry mating system effect on aggression rate. Referred in main text as S5.1.

a)

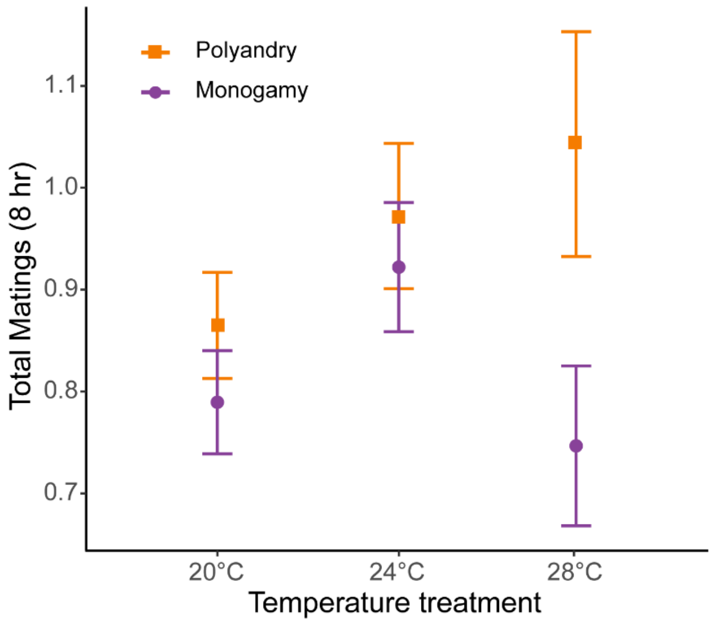


b)

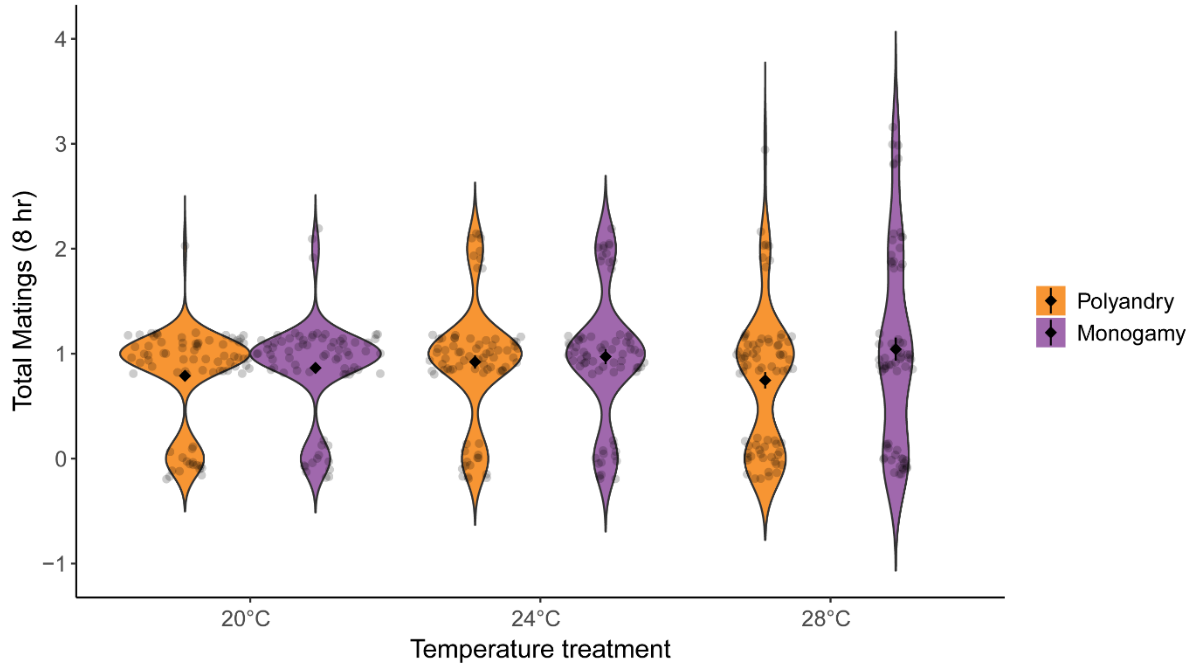


**Figure 5 – figure supplement 2**. a) Total number of matings across the 8 hr of observations. Data from reproductive behaviour measures (mean ± SEM) across temperature and mating system treatments. b) Violin plot. Referred in main text as S5.2.

a)

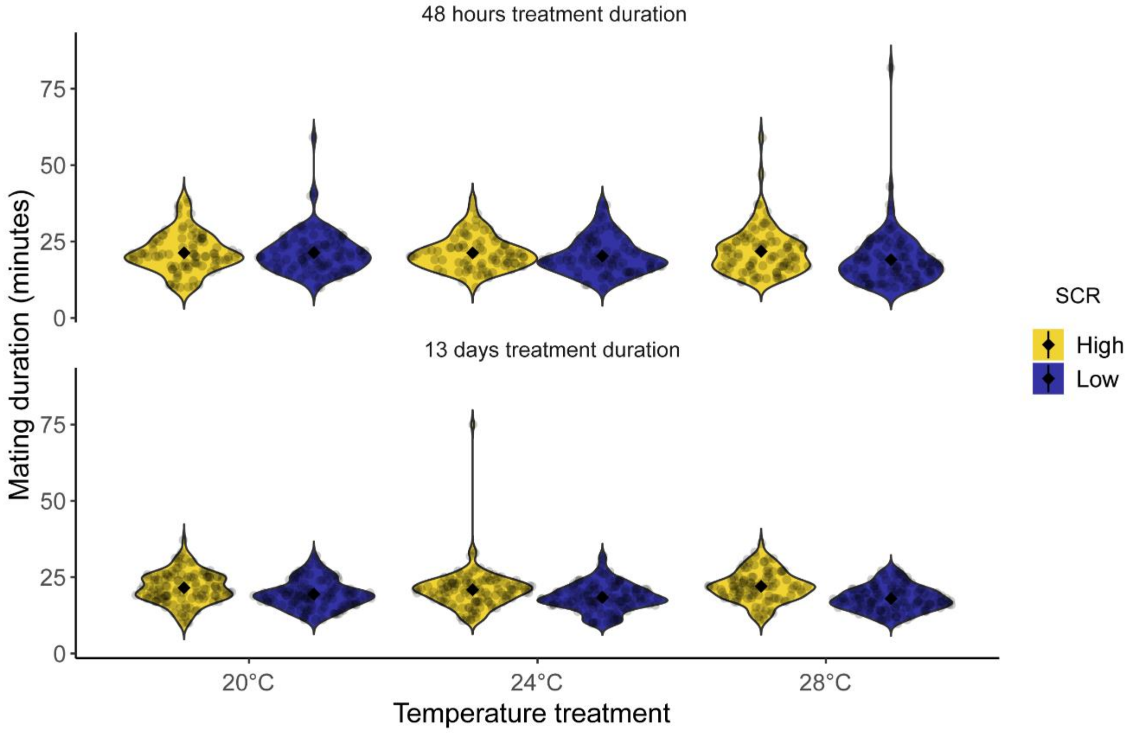


b)

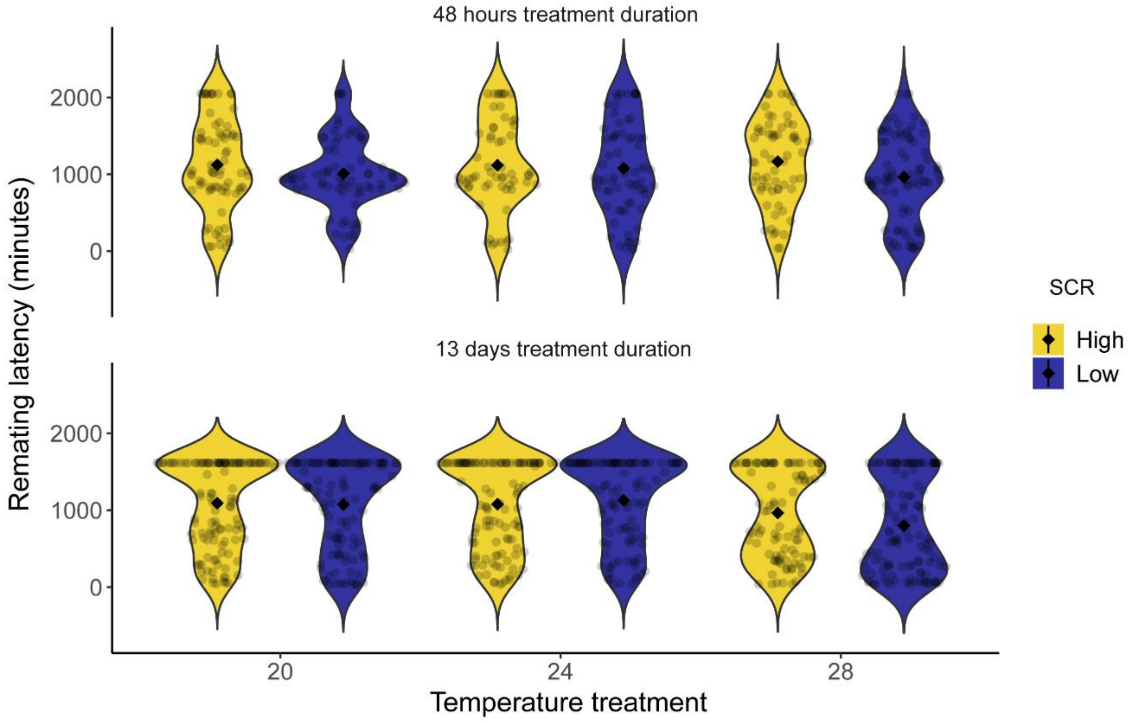


**Figure 6 – figure supplement 1.** Violin plots for a) Mating duration of males exposed to a high (8 males per vial) or low sperm competition risk (1 male per vial) level 48 hours (experiment 2) and 13 days (experiment 3) before mating across temperature treatments, b) female remating latency following a single mating with either a male from a high or low sperm competition risk level, for both 48 hours and 13 days of temperature treatment duration before mating in a common garden. Referred in main text as S6.1.

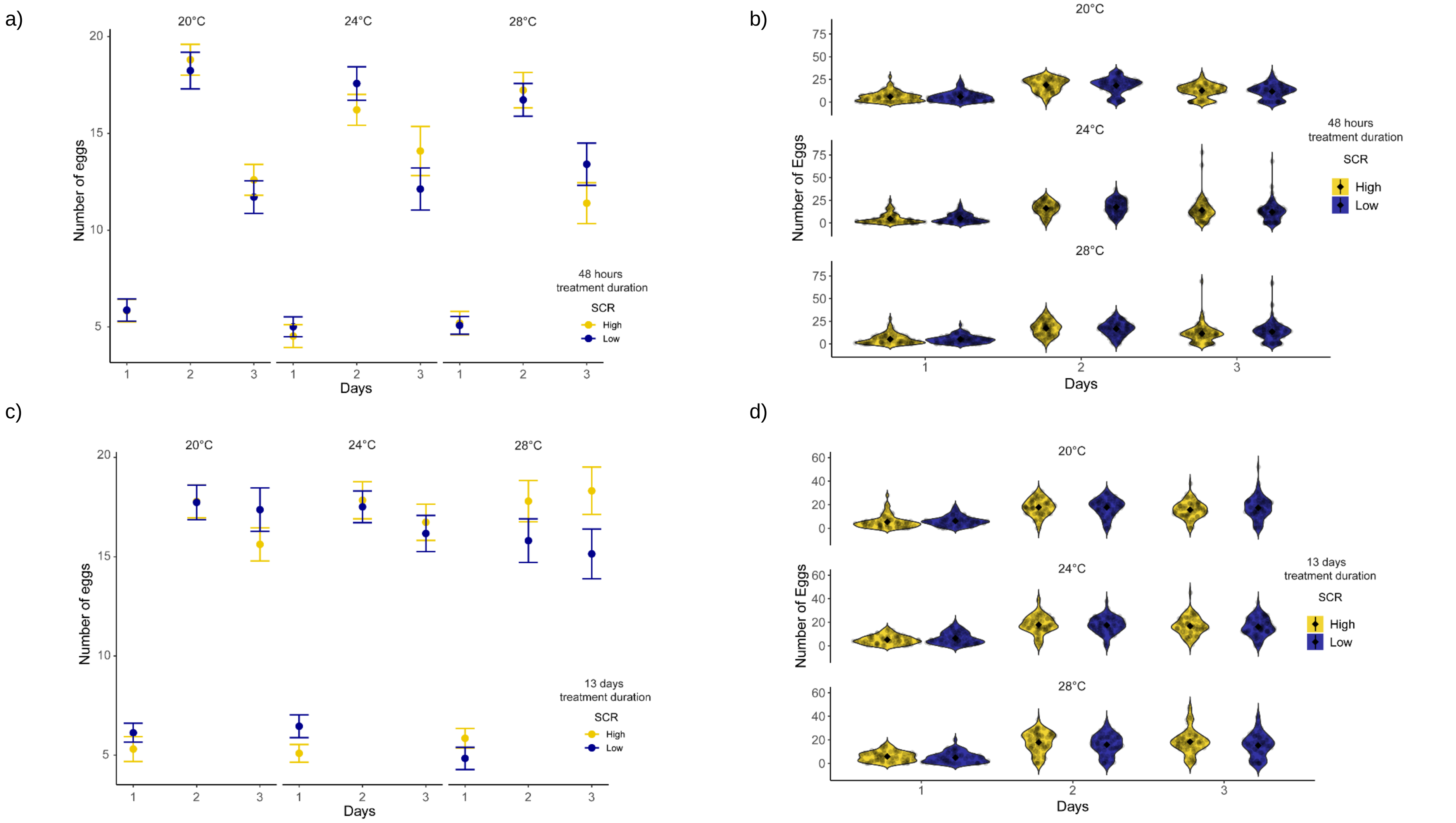


**Figure 6 – figure supplement 2**. Eggs produced by females during the first three days following a single mating with treated males. a) mean ± SEM, b) violin plot, 48 hours treatment duration. c) mean ± SEM, d) violin plot, 13 days treatment duration. Referred in main text as S6.2.

a)

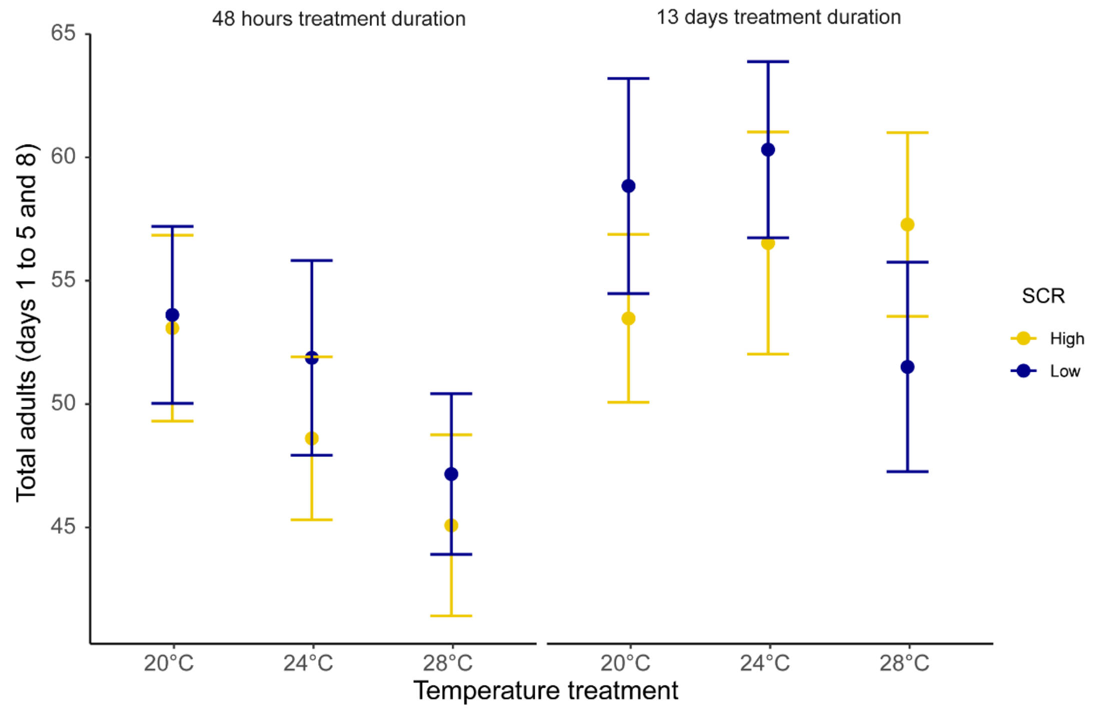


b)

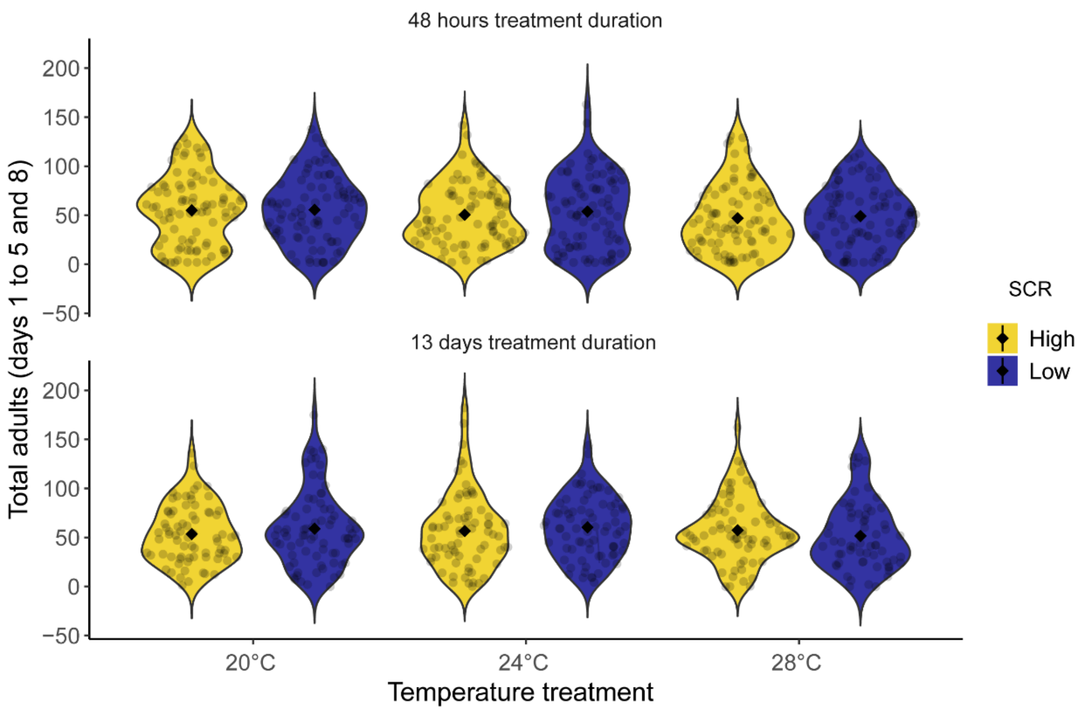


**Figure 6 – figure supplement 3.** a) Total of offspring produced by females during the days 1, 2, 3, 4, 5, and 8 after mating (mean ± SEM) following a single mating with treated males. b) Violin plot. Referred in main text as S6.3.

a)

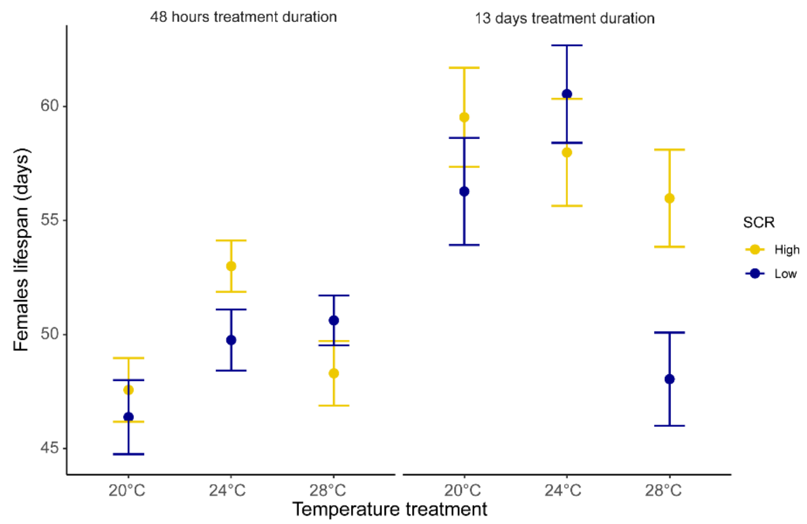


b)

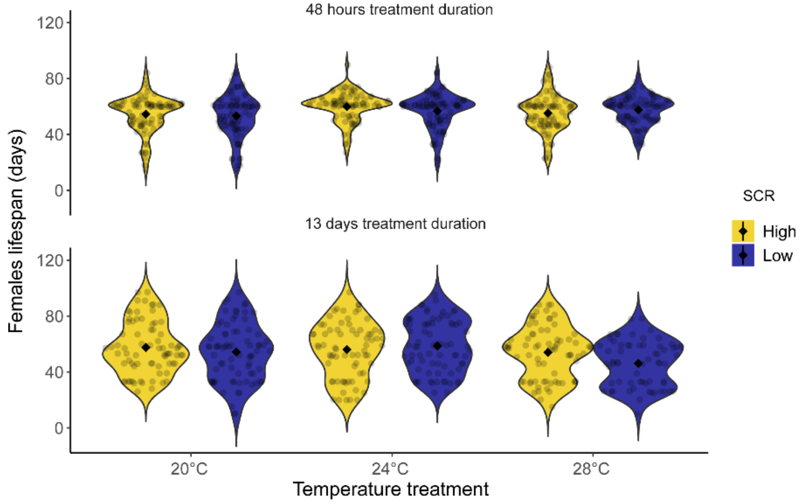


c)

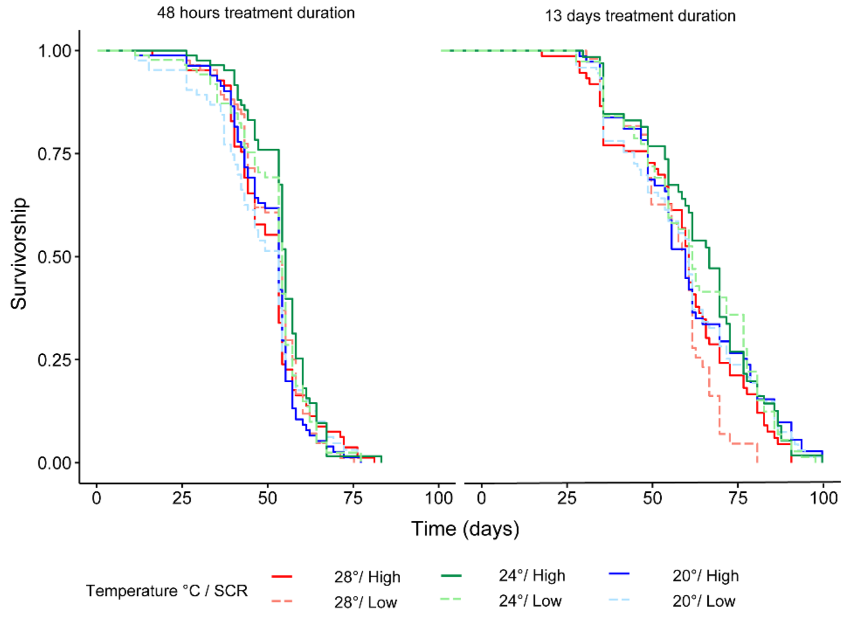


**Figure 6 – figure supplement 4.** a) Female lifespan after mating (mean ± SEM) following a single mating with treated males. b) Violin plot c) Survival plot from the Cox proportional hazard model. Referred in main text as S6.4.

**Table 1 – table supplement 1**. Summary statistics from Tukey's post hoc test to examine the meaning of significant interactions between temperature and mating system effects. a) Polyandry – Monogamy contrast table for each temperature level for female fitness components. b) Polyandry – Monogamy contrast table for each temperature level for underlying behavioural mechanisms. Test from GLMs fitted with temperature as factor. Note that using Tukey's post hoc yielded qualitatively identical results from running models separately for each temperature. Referred in main text as S1.1.

a)

| ***T*°C** | *LRS* | | | | *Reproductive ageing* | | | | *Actuarial ageing* | | | |
|---|---|---|---|---|---|---|---|---|---|---|---|---|
| | *T ratio* | *Df* | *p* | *Estimate +/- SE* | *T ratio* | *Df* | *p* | *Estimate +/- SE* | *T ratio* | *Df* | *p* | *Estimate +/- SE* |
| 20° | 2.31 | 422 | 0.022 | 3.73±1.6 | -2.9 | 422 | 0.004 | -16±5.5 | 8.05 | 425 | <0.001 | 14.9±1.8 |
| 24° | 3.66 | 422 | <0.001 | 5.99±1.6 | -6.19 | 422 | <0.001 | -34.4±5.6 | 5.2 | 425 | <0.001 | 9.69±1.8 |
| 28° | 1.35 | 422 | 0.177 | 2.27±1.7 | -4.16 | 422 | <0.001 | -23.8±5.7 | 3.12 | 425 | 0.002 | 5.95±1.9 |

b)

| ***T*°C** | *Courtship rate* | | | | *Rejection rate* | | | |
|---|---|---|---|---|---|---|---|---|
| | *T ratio* | *Df* | *p* | *Estimate +/- SE* | *T ratio* | *Df* | *p* | *Estimate +/- SE* |
| 20° | 0.14 | 438 | 0.891 | 0.25±1.8 | -0.48 | 438 | 0.628 | -0.07±0.14 |
| 24° | -5.25 | 438 | <0.001 | -9.66±1.8 | -3.6 | 438 | <0.001 | -0.5±0.14 |
| 28° | -5.91 | 438 | <0.001 | -11.0±1.9 | -4.15 | 438 | <0.001 | -0.62±0.15 |

**Table 1 – table supplement 2.** a) Summary statistics from Cox PH survival model as a complementary analysis to analyze potential differences in mortality risk across treatments from the experiment 1. b) Summary statistics from fitting Cox PH models are shown separately for each temperature level due to a significant interaction between temperature and mating system. c) Polyandry – Monogamy contrast table from Tukey's post hoc for each temperature level from Cox PH survival model fitted with temperature as factor. p-values from Cox HP models are computed using ANOVA type III, LR test. Note that using Tukey's post hoc yielded qualitatively identical results from running models separately for each temperature. The corresponding survival plot is plotted in Figure S4.1. Referred in main text as S1.2.

a)

| *Effect* | *Chisq* | *Df* | *p value** |
|---|---|---|---|
| *Temperature * Mating System* | 7.18 | 1 | 0,007 |
| *Mating System* | 16.14 | 1 | <0,001 |
| *Temperature* | 62.60 | 1 | <0.001 |

* p-values were corrected for multiple testing using BH correction

b)

| *T°* | *Chisq* | *Df* | *p value** | *Estimate (95% CI)* |
|---|---|---|---|---|
| 20°C | 43.42 | 1 | <0.001 | 1.87 (1.55 - 2.26) |
| 24°C | 40.06 | 1 | <0.001 | 1.83 (1.52 - 2.22) |
| 28°C | 18.21 | 1 | <0.001 | 1.46 (1.23 - 1.74) |

* p-values were corrected for multiple testing using BH correction

c)

| *T°* | *Estimate* | *SE* | *Df* | *T ratio* | *p value* |
|---|---|---|---|---|---|
| 20°C | 1.27 | 0.18 | Inf | 6.98 | <0.001 |
| 24°C | 1.29 | 0.17 | Inf | 7.36 | <0.001 |
| 28°C | 0.59 | 0.17 | Inf | 3.47 | <0.001 |

**Table 3 – table supplement 1**. Summary statistics from Tukey's post hoc test as a complementary analysis to examine the meaning of significant interactions found for mating duration and remating latency. a) High – low sperm competition risk contrast table for each temperature level. b) Long – short treatment duration contrast table for each temperature level. c) High – low sperm competition risk contrast table for each treatment duration. Test from GLMs fitted with temperature as factor. Note that using Tukey's post hoc yielded qualitatively identical results from running models separately for each temperature or treatment duration. Referred in main text as S3.1.

a)

| ***T*°C** | *Mating duration** | | | | *Remating latency** | | | |
|---|---|---|---|---|---|---|---|---|
| | *T ratio* | *Df* | *p* | *Estimate +/- SE* | *T ratio* | *Df* | *p* | *Estimate +/- SE* |
| 20° | 1.7 | 1232 | 0.08 | 0.05±0.02 | 1.109 | 1087 | 0.268 | 66.7±60.1 |
| 24° | 2.98 | 1232 | 0.002 | 0.08±0.02 | -0.071 | 1087 | 0.943 | -4.5±62.6 |
| 28° | 5.74 | 1232 | <0.001 | 0.17±0.02 | 2.969 | 1087 | 0.003 | 183.2±61.7 |

*Results are averaged over the levels of treatment duration

b)

| ***T*°C** | *Mating duration** | | | | *Remating latency** | | | |
|---|---|---|---|---|---|---|---|---|
| | *T ratio* | *Df* | *p* | *Estimate +/- SE* | *T ratio* | *Df* | *p* | *Estimate +/- SE* |
| 20° | -1.4 | 1232 | 0.143 | -0.04±0.03 | -0.022 | 1087 | 0.982 | -1.32±60.1 |
| 24° | -1.9 | 1232 | 0.046 | -0.05±0.03 | -0.216 | 1087 | 0.828 | -13.5±62.6 |
| 28° | -0.8 | 1232 | 0.41 | -0.02±0.03 | -3.259 | 1087 | 0.001 | -201.1±61.7 |

*Results are averaged over the levels of sperm competition risk

c)

| ***Treatment duration*** | *Mating duration** | | | |
|---|---|---|---|---|
| | *T ratio* | *Df* | *p* | *Estimate +/- SE* |
| Short (48 hours) | 2.41 | 1232 | 0.016 | 0.06±0.02 |
| Long (13 days) | 6.33 | 1232 | <0.001 | 0.143±0.02 |

*Results are averaged over the levels of temperature

**Table 4 – table supplement 1.** Summary statistics from the Hurdle model to analyze potential differences in egg production across treatments with temperature as a factor. Note that using temperature as a factor yielded qualitatively identical results than treating it as a continuous covariable. p-values from Hurdel model are computed using ANOVA type III, Wald test. Corresponding data is plotted in Figure S6.2. Referred in main text as S4.1.

| *Effect* | *Chisq* | *Df* | *p value* |
|---|---|---|---|
| *SCR* | 0.0016 | 1 | 0.968 |
| *Temperature* | 2.7419 | 2 | 0.253 |
| *Treatment duration* | 9.0076 | 1 | 0.002 |
| *SCR*Treatment duration* | 0.039 | 1 | 0.843 |
| *SCR*Temperature* | 1.9116 | 2 | 0.384 |
| *Temperature*Treatment duration* | 2.7871 | 2 | 0.248 |

**Table 4 – table supplement 2**. Summary statistics from Tukey's post hoc test as a complementary analysis to examine the meaning of significant interaction between temperature and treatment duration for total of offspring produced by females during the days 1, 2, 3, 4, 5, and 8 after mating. Short (48 hours) – Long (13 days) treatment duration contrast table for each temperature level. Test from GLMs fitted with temperature as factor. Note that using Tukey's post hoc yielded qualitatively identical results from running models separately for each temperature. Referred in main text as S4.2.

| ***T*°C** | *Total of adults* | | | |
|---|---|---|---|---|
| | *T ratio* | *Df* | *p* | *Estimate +/- SE* |
| 20° | 0.74 | 947 | 0.453 | 2.81±3.76 |
| 24° | 2.17 | 947 | **0.029** | 8.19±3.76 |
| 28° | 2.18 | 947 | **0.028** | 8.44±3.86 |
